# Autism spectrum disorders: an impaired glycolysis induces an ATP deficiency and a reduced cell respiration

**DOI:** 10.1101/2024.01.22.576644

**Authors:** Francois Féron, Damien Caillol, Laure Fourel, Silke Leimkuhler, Olga Iranzo, Bruno Gepner, Gaelle Guiraudie-Capraz

## Abstract

In two previous studies, based on human olfactory ecto-mesenchymal stem cells (OE-MSC) of 11 patients with autism spectrum disorders (ASD) and 11 healthy individuals, we demonstrated that the lower abundance of the enzyme MOCOS (MOlybdenum COfactor Sulfurase) and its associated lower expression of the long non-coding RNA, COSMOC, induces neurotransmission and synaptic defects as well as an exacerbated oxidative stress sensitivity. To move a step further, we assessed whether these defects were associated to a disturbed mitochondrial homeostasis. For that purpose, we used cellular and molecular techniques to quantitfy mitochondrial metabolism and biogenesis, ATP production and cell respiration in OE-MSCs from the 8 ASD patients of the cohort that display the most severe symptoms. We show here that OE-MSCs from ASD patients, when compared to control individuals, display i) a reduced expression/abundance of glycolysis-associated transcripts and metabolites, ii) an overall reduced ATP, mainly due to the impaired glycolysis, iii) a reduced basal cell respiration and iv) a modified mitochondrial network. These results are in accordance with some of our previously published data and may explain some of the symptoms – stress, overarousal, seizures, increased or decreased muscle tone, fatigue - observed in autism spectrum disorders.

## Introduction

Autism Spectrum Disorders (ASD), which affect around 50 million individuals worldwide (Hahler and Elsabbagh 2015), is a major public health issue. With an onset under the age of three, ASD are understood as diseases arising from pre- and/or early post-natal brain developmental anomalies and/or early brain insults/infections. To unveil the molecular mechanisms at play during the misshaping of the developing brain, we chose to study cells representative of the very early stages of ontogeny, namely adult stem cells and, more specifically, human nasal olfactory stem cells (Delorme et al. 2010). In a first series of experiments, based on a cohort of 11 autistic patients and 11 healthy individuals we demonstrated the reduced expression/production of *MOCOS* and MOCOS (MOlybdenum COfactor Sulfurase), an enzyme involved in purine metabolism^3^, and its associated long non-coding RNA that we named COSMOC^4^. We observed that reduced expression of MOCOS and/or COSMOC induces neurotransmission and synaptic defects as well as an exacerbated oxidative stress sensitivity (Féron et al. 2016; Rontani et al. 2021).

All these molecules are involved in the synthesis of molybdenum enzymes, which begins in mitochondria with the conversion of guanosine triphosphate (GTP) to cyclic pyranopterin monophosphate (cPMP) by the two active forms of the MOCS1 enzyme (MOCS1A and MOCS1AB) (Mendel and Kruse 2012). cPMP is then converted outside the mitochondria *via* MOCS2 to molybdopterin (MPT), which in turn is further converted to the MOlybdenum COfactor (MOCO) by gephyrin (GPHN). MOCO is the central element of the four human molybdenum enzymes: i) sulfite oxidase (SUOX) that transforms sulfites into sulfates (Mendel and Schwarz 2023); ii) xanthine dehydrogenase (XDH) and aldehyde oxidase (AOX) that catalyse the two-step reaction from hypoxanthine to xanthine and from xanthine to uric acid in purine metabolism (Bortolotti et al. 2021); iii), and is involved in the metabolism of retinaldehyde, respectively, and the mitochondrial amidoxime-reducing component (mARC), a key factor in the N-hydroxylation of prodrugsor N-oxygenated compounds (Struwe, Scheidig, and Clement 2023).

Many observations have linked autism disorders to mitochondrial dysfunction (for a review, (Nabi et al. 2023)). An exploratory study reported that the majority of lymphocytes from very young ASD children exhibit a reduced activity of oxidative phosphorylation as well as a mitochondrial DNA over-replication (Giulivi et al. 2010). In addition to mutations in mitochondrial DNA (Levinger, Giegé, and Florentz 2003; Weissman et al. 2008; Napoli et al. 2018; HADJIXENOFONTOS et al. 2013), recurrent studies found anomalies in i) electron transport chain (ETC) complex activity (Haas 2010; Boddaert et al. 2009; Gu et al. 2013; Anitha, Nakamura, Thanseem, Yamada, et al. 2012; Anitha, Nakamura, Thanseem, Matsuzaki, et al. 2012), ii) Krebs cycle metabolites (Rossignol and Frye 2012; Suomalainen et al. 2011; Weissman et al. 2008; Filipek et al. 2003) and iii) oxidative stress and redox regulation (Legido 2018; Adams et al. 2009; Borrás et al. 2003). However, none of these studies was able to make causal inferences between disturbed mitochondrial metabolism and ASD. To move a step further, we used a panel of cellular and molecular techniques to unveil mitochondrial disturbances in stem cells originating from a peripheral nervous tissue, namely the olfactory mucosa, of 8 ASD patients with severe symptoms and 8 age- and gender-matched healthy individuals.

## Materials and methods

### Culture of human olfactory ecto-mesenchymal stem cells

The current study is a follow up of two previous works (Féron et al. 2016; Rontani et al. 2021) based on biopsies from healthy donors and ASD patients. Procedures were approved by the ethical committee (Comité de Protection des Personnes, files #205016 and #205017). All individuals involved in this study provided informed consent in accordance with the Declaration of Helsinki and French laws relating to biomedical research (Féron et al. 2016; Rontani et al. 2021). For the current study, we chose to select ASD patients with the most severe symptoms, i.e. individuals classified as level 3. According to the DSM-5, those at this level need a great deal of support. In our cohort, 8 of the 11 individuals fall into this category. Olfactory ecto-mesenchymal stem cells (OE-MSC) were cultivated in proliferating conditions, using DMEM/HAM F12 culture medium (Life Technologies) supplemented with 10% fetal calf serum (Life Technologies) and penicillin/streptomycin (1%) (Life Technologies).

### Metabolome analysis

OE-MSCs from 8 ASD patients and 6 control subjects were incubated with 400 µL of cold methanol (−20°C) for 30 minutes, before centrifugation and filtration to remove residual proteins from the samples. After drying under a stream of nitrogen, the dry extract was taken up in 250 µL of a water/acetonitrile mixture (9/1, v/v). Samples were separated using UPLC Ultimate 3000 (Thermo Scientific), coupled to a Q-Exactive Plus quadrupole orbitrap hybrid high-resolution mass spectrometer (HRMS) equipped with an electrospray ionisation source (H-ESI Il). Chromatographic separation was carried out on a binary solvent system using two columns, with specific conditions for each. The mass spectrometer was calibrated in positive and negative mode using two calibration solutions and a solution composed of caffeine, n-butylamine, MRFA (peptide) and Ultramark 1621. Calibration in negative mode was carried out using a solution composed of sodium dodecyl sulphate, a solution of taurocholate and Ultramark 1621. The molecules separated by chromatography were analysed in positive and negative ionisation mode in the same cycle. Mass spectra were recorded using a resolution of 35,000 FWHM for a theoretical mass-to-charge ratio (m/z) of 200. To ensure good analytical repeatability, quality control (QC) samples were prepared by taking an equal amount from each biological sample.

From the list of 187 metabolites identified, two ratios – A/B and 1/(A/B) – were calculated where A corresponds to the quantity of metabolite X for every ASD patient and B corresponds to the mean quantity of the same metabolite in the control population. The cut-off value for fold change was set at ±1,5. The first ratio identifies any higher or lower abundance of metabolites in every patient while the second ratio reflects the percentage associated with these changed concentrations. Only metabolites with an altered abundance in at least 4 ASD patients were used for further analysis.

### Transcriptome analysis

Each RNA sample was quantified on Nanodrop and quality was examined on the Fragment Analyzer (Agilent). 2µg of total RNA from 17 samples of human (RIN >9) were used as the starting material in the TruSeq Stranded mRNA Illumina kit according to the manufacturer’s instructions. Each library was barcoded using TruSeq_RNA_LT Indexes (Illumina), according to manufacturer’s instructions (A001 to A017).

Each library was quantified on Qubit with Qubit® dsDNA HS Assay Kit (Life Technologies) and then, size distribution was examined on the Fragment Analyzer with High Sensitivity NGS Fragment Analysis kit (Agilent), to ensure that the samples were the proper size, no adaptor contamination, and to estimate the sample molarity. Each library was diluted to 4 nM and then pulled together at equimolar ratio. The denaturation was performed with 5 µL of pooled libraries (4 nM) and 5 min incubation with 5 µL of fresh NaOH (0.2N) and then addition of 5 µL of fresh Tris-HCl (200 mM - pH 7), according to manufacturer’s instructions.

The dilution of 20 pM pooled libraries was performed with HT1 to a 1.6 pM final concentration. PhiX library as a 1% spike-in for use as a sequencing control was denatured and diluted, according to manufacturer’s instructions, and 1,2 µL was added to denatured and diluted pooled libraries before loading. Finally, libraries were sequenced on a High-output flow cell (400M clusters) using the NextSeq® 500/550 High Output 150 cycles kit (Illumina), in paired-end 76/76 nt mode, according to manufacturer’s instructions.

### ATP quantification

Cellular bioenergetics were measured using the Seahorse XF Cell Mito Stress Test kit, which quantifies the oxygen consumption and ATP production, as previously described by the team (Ogunkola et al. 2023). Briefly, OE-MSCs were plated, at the density of 15,000 c/well, on 24-well seahorse plates in DMEM/HAM medium a night prior to analysis. Oxygen consumption rate (OCR) after addition of oligomycin (1 µM) was used to estimate the rate of ATP production. Carbonyl cyanide-4-trifluoromethoxy phenylhydrazone (FCCP, 0.5 µM) was added to assess the maximal mitochondrial respiratory capacity. The flux of electrons through complex III and I was blocked with antimycin A (0.5 µM) and rotenone (0.5 µM), respectively. Any residual activity in the presence of these inhibitors was considered as non-mitochondrial OCR. Results were normalized to the number of cells, using a high content screening microscope (Operetta, CLS, Perkin Elmer).

### Fluorescent labelling of mitochondria

OE-MSCs from control subjects and ASD patients, seeded on glass coverslips, were incubated for 15 minutes at 37°C with Mitotracker™ Red CMXRos (50nM, ThermoFisherScientific) and 1:500 Hoechst blue (ThermoFisher). After rinsing with PBS, cells were fixed with 4% paraformaldehyde for 10 minutes and rinsed three times with PBS. The coverslips were then mounted on slides with a glue containing ProLong™Gold antifade reagent (ThermoFisher), which limits the loss of fluorescence. Mitochondrial markings were visualised and digitised using confocal microscope (inverted Zeiss Axio Observer Z1 LSM700) equipped with a ×63 (NA 1.4) oil, Plan-Apochromat objective (Zeiss).

### Analysis of mitochondrial network

The confocal images needing to be improved for increased sharpness, an image processing was adapted from the work of Valente et al. 2017 and Merrill et al. 2017 (Valente et al. 2017; Merrill, Flippo, and Strack 2017). They were transformed into binary images that were analysed using the “Mitochondrial Morphology” macro-keyboard of Fiji software (https://fiji.sc/), which measures, among other things, mitochondrial interconnectivity and elongation. From the binary image, the ImageJ software allowed to obtain an image of the mitochondrial skeleton, analysed by the ‘Analyze Skeleton’ macro-keyboard, which counts the different mitochondrial elements and branches, with their average and maximum size. Mitochondria were classified according to their organisation, shape and size. Organised in a network or isolated (individual mitochondria), three forms can be distinguished: punctate mitochondria, large/round mitochondria and rod mitochondria.

### Western blot of proteins involved in fusion/fission of mitochondrial network

Proteins were extracted from 12 OE-MSC pellets (n = 6 ASD patients and 6 control subjects). Cells were mechanically lysed using a solution composed of RIPA buffer (Sigma-Aldrich) and 1/100 protease inhibitors (Calbiochem). After 30 minutes of incubation at 4°C, followed by sonication (2 minutes), protein concentrations were determined by a Lowry assay (DC™ Protein Assay, BioRad) and protein samples were stored at −20°C until use. For SDS-PAGE electrophoreses, 50 μg of protein samples were denatured in 2X Laemmli buffer (Bio Rad) and heated at 95°C for 5 minutes. Denatured protein samples were separated on a 12% SDS-PAGE polyacrylamide gel for 120 minutes at a constant 0.07 Ampere per gel. Proteins were transferred with the PowerBlotter XL (Invitrogen by ThermoFisher Scientific) onto nitrocellulose membranes (Amersham™ Protran™ Premium 0.2μm, GE Healthcare), using the Standard Semidry protocol (25 V constant, 0.2A for 60 minutes) with transfer buffer (0.25 M Tris, 1.92M glycine, 200 mL methanol and 700 mL osmosed water at pH 8.3). For immunodetection, the non-specific sites were saturated by incubation at room temperature with agitation for 90 minutes in a solution of 1X PBS/5% milk/0.2% Tween (TWEEN©20, Sigma-Aldrich). The membranes were then incubated overnight at 4°C in a humidity chamber with the primary antibody diluted in PBS 1X/milk 5%/Tween 0.2%. The following day, after 3 washes with PBS 1X/Tween 0.2%, the membranes were incubated with the peroxidase-coupled secondary antibody (HRP) diluted in PBS 1X/5% milk/Tween 0.2%, at room temperature, under agitation for 80 minutes. After washing with PBS 1X/Tween0.2%, the proteins of interest were revealed by chemiluminescence using the ECL WesternBlot detector kit (GE Healthcare) and the results were digitised in a UVITEC11Photo Imager (Ubitec Photoimager, Dutscher) using Nine Alliance software (Alliance©Software, UVITEC). For relative quantification, the optical density (OD) of each immunodetected protein band was measured using Fiji software (National Institutes of Health - https://imagej.net/Fiji) and normalised to the OD of the GAPDH band.

### Statistics

All statistics are presented as mean ± 1 standard error of the mean (SEM). All distributions were tested for normality (Shapiro-Wilk test, significant at 0.05) and homogeneity of variance (Fisher test, significant at 0.05) prior to analysis. Based on the results of these initial tests, the comparisons were carried out either with Student’s t-test (normal distribution and homogeneity of variances) or its variant Welch corrected t-test (normal distribution and heterogeneity of variances) or with the Mann-Whitney test (non-normal distribution). The significance scores are: * for p < 0.05, ** for p < 0.01 and *** for p < 0.001. A difference was considered significant if the p-value was less than 0.05. All statistical analyses were carried out using GraphPad Prism software version 8.4.0 for OS X, GraphPad Software, San Diego, California, USA, www.graphpad.com.

## Results

### The abundance of many mitochondria-associated metabolites is disrupted in ASD stem cells

Quantification of metabolites between each ASD patient and the mean of 6 control individuals reveals an altered abundance of molecules involved in four mitochondria-related pathways (figure 1A). For the current analysis, the cut-off value for fold change was set at ±1,5. Among the most striking findings is the lower abundance of adenine and pyruvate in all patients. The lower abundance of the former, together with a lower abundance of adenosine diphosphate (ADP) in 6 out 8 patients, indicate a putative disruption of oxidative phosphorylation. Altered abundance of the latter, combined with a reduced production of glucose-6-phosphate (6/8 patients) and ascorbic acid (7/8), point to a presumed modified glycolysis. It should be noted that ketoglutarate, a molecule involved in the Krebs cycle as well as in carnitine biosynthesis, is present in lower concentration in 7 ASD patients. All the abnormalities in metabolites associated with mitochondria are summarized schematically in Figure 1B.

**Figure 1.**
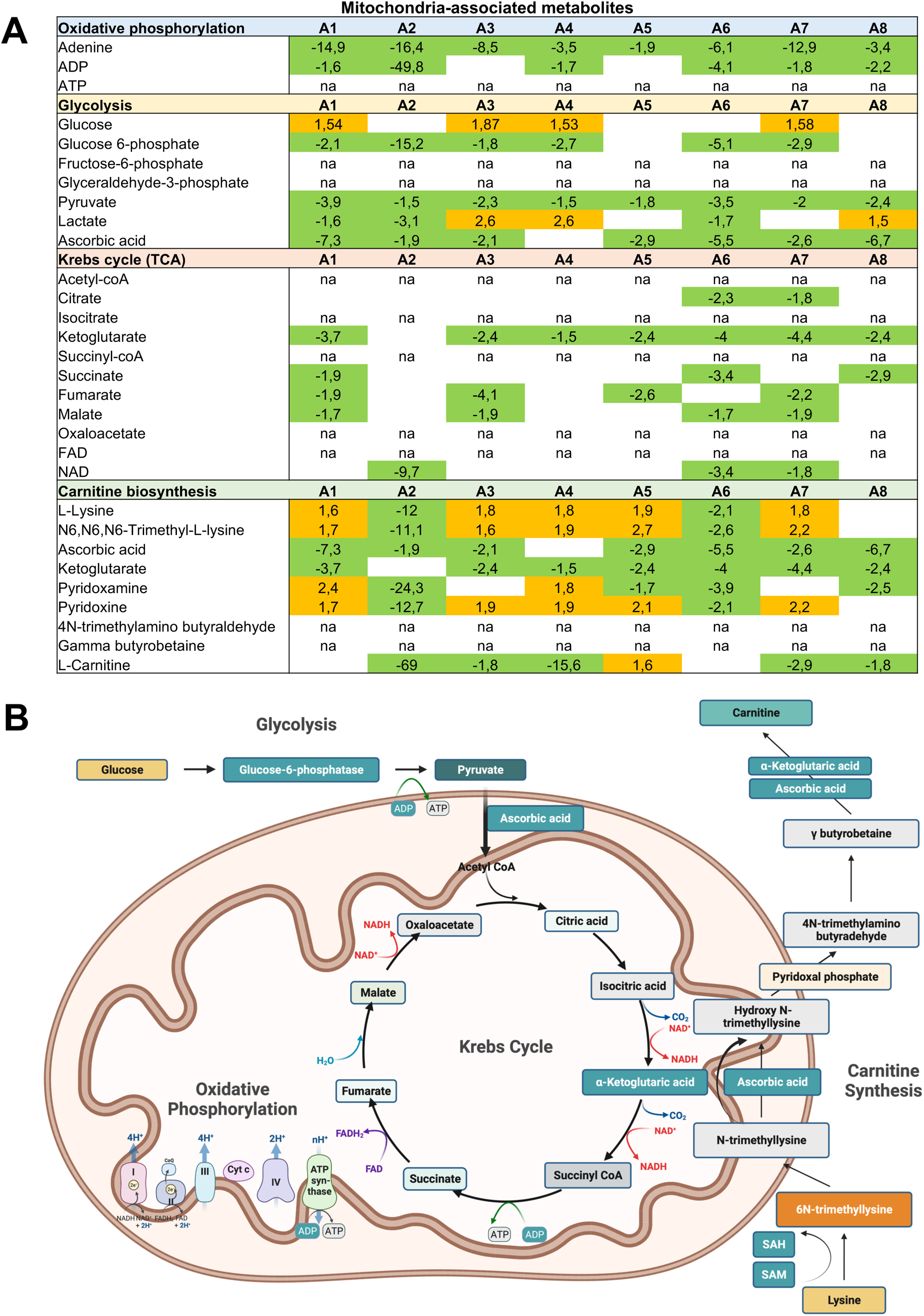
Mitochondria-associated metabolome. A) Main results for the 4 metabolic occurring, wholly or partially, in the mitochondria of olfactory stem cells. The cut-off value for fold change was set at ±1,5. To varying degrees, all pathways are affected in cells from ASD patients, compared with mean values from control individuals. B) Schematic summary of the main findings. The tone of each color correlates with the number of concerned patients. Green: underabundance; orange: overabundance; na: not available.

### ASD stem cells show reduced ATP production

The widespread deficiency of adenine and, to a lesser extent, ADP led us to consider a disturbance in oxidative phosphorylation. To make sure it was the case, we studied ATP production using the Seahorse apparatus and its various associated protocols. Figure 2A shows the different values measured with the addition of specific inhibitors at various stages of this metabolic pathway, which includes 5 complexes. The difference between the two arms of the cohort is illustrated in Figure 2B, which describes the tracings of a control individual and two ASD patients. Before examining the data for each individual in more detail, it is essential to note that olfactory stem cells, which are highly proliferative cells just like cancer cells, differ from other somatic cells in that they produce ATP mainly from glycolysis (70%) and not from oxidative phosphorylation (Figure 2C). Overall, ASD patients differ from the control population by a reduction in total ATP production (Figure 2D). This is primarily due to a very significant decrease in glycolysis-associated ATP production (Figure 2E), which is not fully compensated for by an increase in ATP produced during oxidative phosphorylation (Figure 2F).

**Figure 2.**
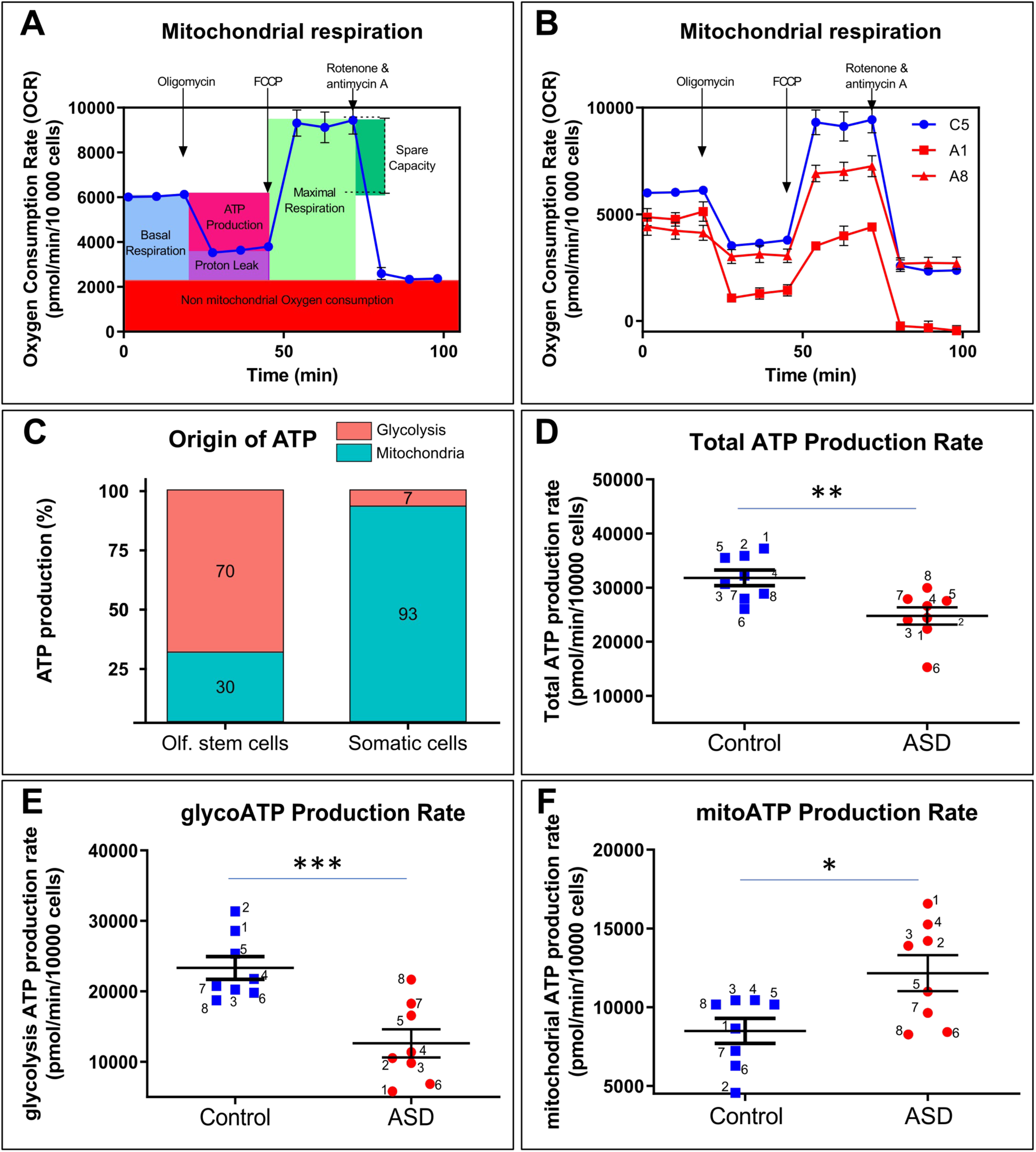
ATP production. A) Schematic representation of the protocol performed with the Seahorse device. B) Examples of measurements of mitochondrial respiration in stems cells from a control individual (blue line) and two ASD patients (red lines). C) Illustration of the relative percentage of mitochondrial ATP and glycolytic ATP in somatic cells and olfactory stem cells. D-F) Box plots summarizing the total ATP, glycoATP and mitoATP levels. * = p<0.05; ** = p<0.01 ; *** = p<0.005

### Many glycolysis-related transcripts are present in lower concentrations in ASD stem cells

The unexpected decrease in ATP production from a reduced glycolysis pathway led us to investigate the causes of this lower abundance. To do this, we carried out a transcriptome study based on RNA sequencing. We first analyzed the expression of transcripts involved in glycolysis. As with the metabolome, we set the cut-off value for fold change at ±1.5. Figure 3A shows that the concentration of 6 of the 9 genes involved in the conversion of glucose into pyruvate is disrupted in at least 2 patients. Special mention should be made of enolase 2, which concentration is lower in 7 ASD patients and higher in the eighth. It is also important to note that the concentration of several glucose and pyruvate transporters is also affected, in variable proportions and sometimes in contradictory directions. All the abnormalities in glycolysis-associated transcripts are summarized schematically in Figure 3B.

**Figure 3.**
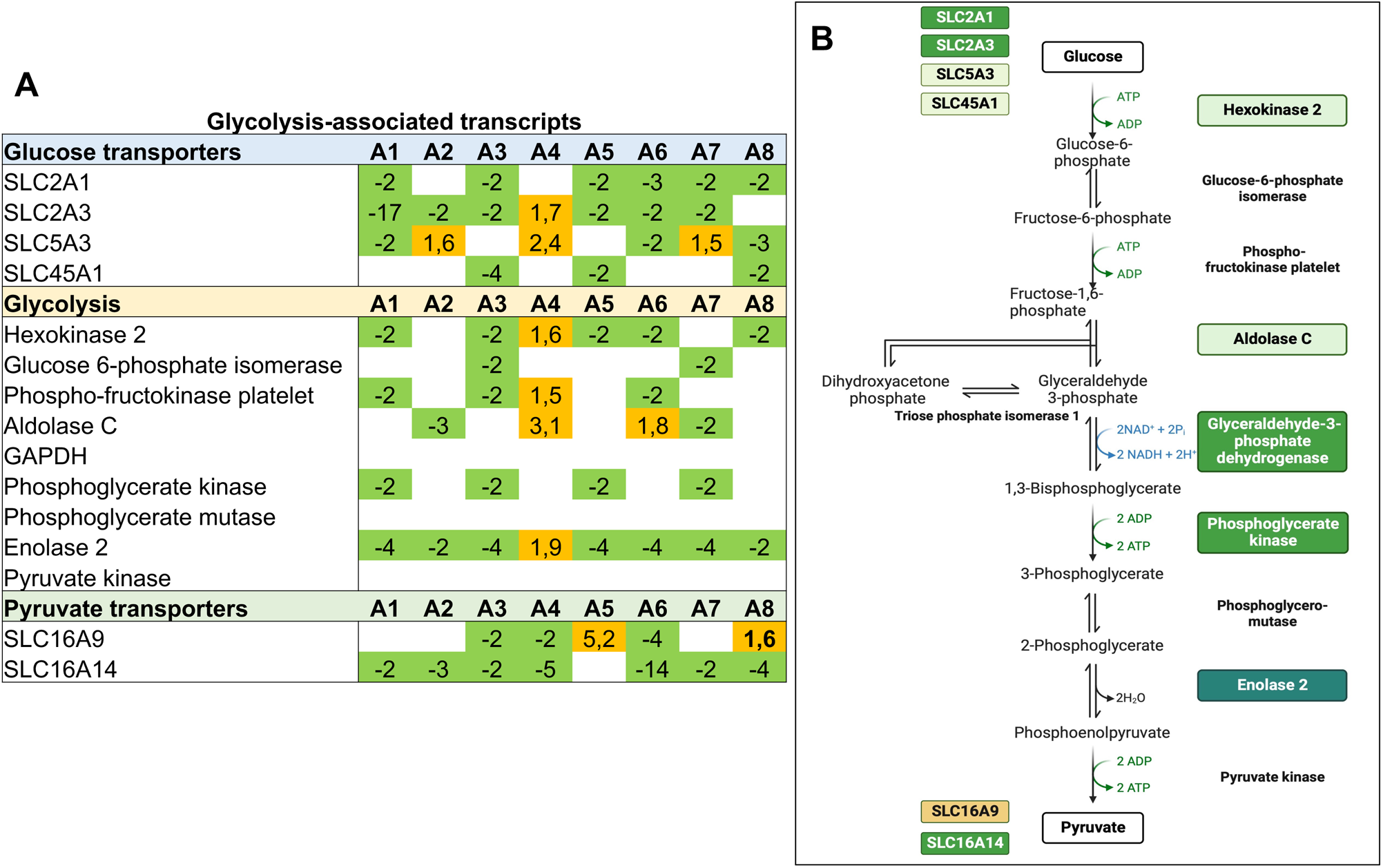
Altered expression of glycolysis-associated transcripts in olfactory stem cells. A) Results for all transcripts involved in glycolysis. Each ASD patient was compared to the mean values of control individuals. B) Schematic view of the altered expression of transcripts encoding proteins involved in glycolysis. The cut-off value for fold change was set at ±1,5. The tone of each color correlates with the number of concerned patients. Green: under-expression; orange: over-expression.

### Stem cells from ASD patients partially rescue ATP production by generating new mitochondria

The metabolome and transcriptome data explain the lower concentration of ATP produced from glycolysis. However, it remains to be explained how stem cells from ASD patients manage to increase the number of ATP molecules derived from oxidative phosphorylation. To understand the mechanisms at work, we incubated the stem cells with a mitochondria-specific fluorescent probe and then analyzed the mitochondrial network. Images 4A and 4B are illustrations of the networks visualized in a control individual and an autistic patient. A detailed study of these networks shows an increase in mitochondrial area (Figure 4C), mitochondrial content (Figure 4D) and the number of punctate mitochondria (Figure 4E) in autistic patients. There was also a trend towards an increase in the number of mitochondria but this did not reach statistical significance (Figure 4F). In order to understand whether this increase in punctate mitochondria is the result of increased fission (Figure 4G) at the expense of reduced fusion (Figure 4J), we measured by Western blot the expression of two markers - DRP1 and OPA1 - specific to mitochondrial dynamics. As shown in Figures 4H and 4I, we observe an increase in the expression of DRP1, a marker of mitochondrial fission. Conversely, Figures 4K and 4L show an under-expression of OPA1, a marker of mitochondrial fusion.

**Figure 4.**
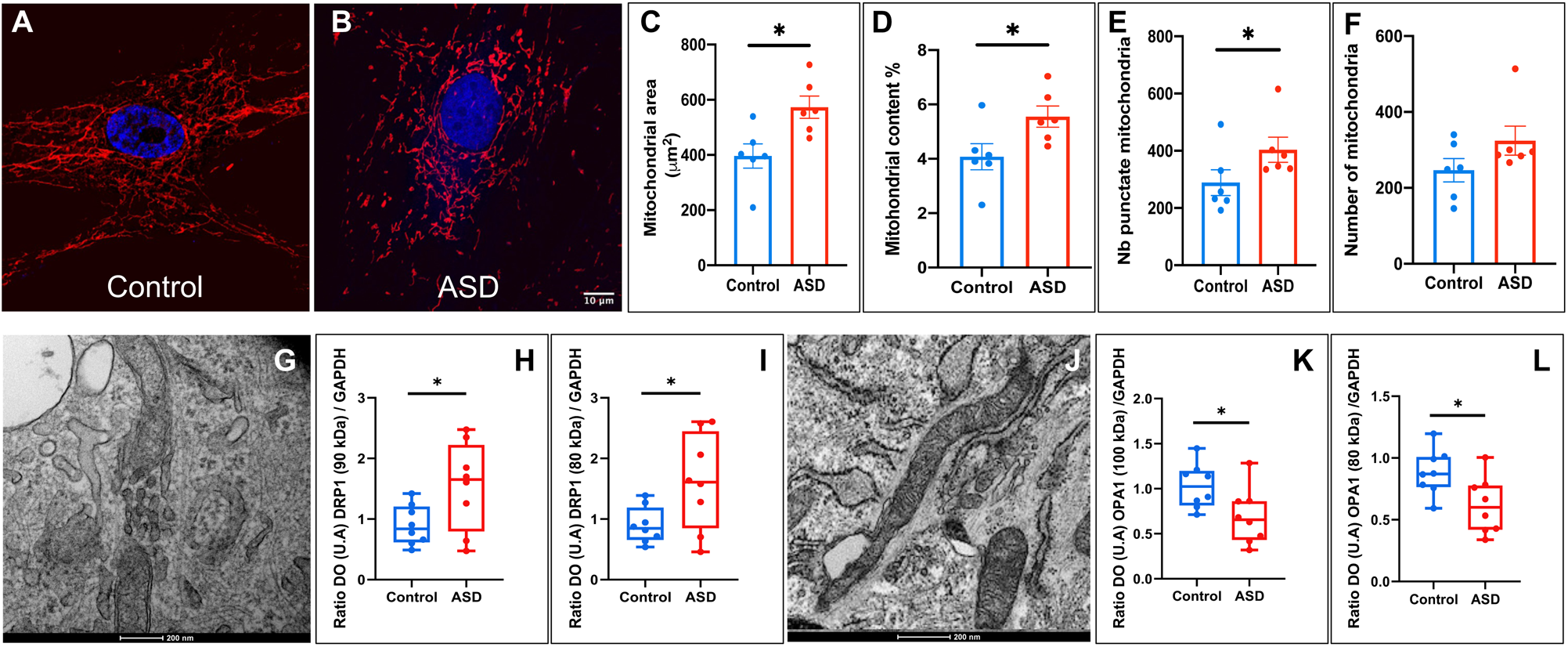
Dynamics of mitochondria. A-B) Examples of the mitochondria network labelled with the mitotracker fluorochrome, in control individual (A) and ASD patient (B), scale bar 10 μm. C-F) The ASD arm of the study shows a statistically significant increase in mitochondrial area (C), mitochondrial content (D), number of punctate mitochondria (E) but not number of mitochondria (F). G) Electron microscopy image of mitochondrial fission, scale bar 200 nm. H-I) ASD stem cells express more abundantly the DRP1 protein involved in mitochondrial fission. J) Electron microscopy image of fused mitochondria, scale bar 200 nm. K-L) ASD stem cells express less abundantly the OPA1 protein involved in mitochondrial fusion. * = p<0.05

### Stem cells from ASD patients exhibit a reduced basal respiration and proton leak

As mentioned above, changes in the mitochondrial network are associated with impaired ATP production. We then assessed whether it was also related to abnormalities in cellular respiration. Figure 5 shows the alterations observed in some ASD patients. Stem cells of several patients (3, 7 and 8) exhibit very low basal respiration and the entire ASD arm of the cohort display a statistically significant difference compared with the control arm (Figure 5A). However, maximal breathing did not differ between the two groups, nor did spare respiratory capacity (Figures 5B,C). Similarly, despite a downward trend, there is no difference between the two groups in ATP-production coupled respiration (Figure 5E). On the other hand, the TSA group differed from the control group by a significant decrease in proton leak (Figure 5D) and a significant increase in coupling efficiency (Figure 5F).

**Figure 5.**
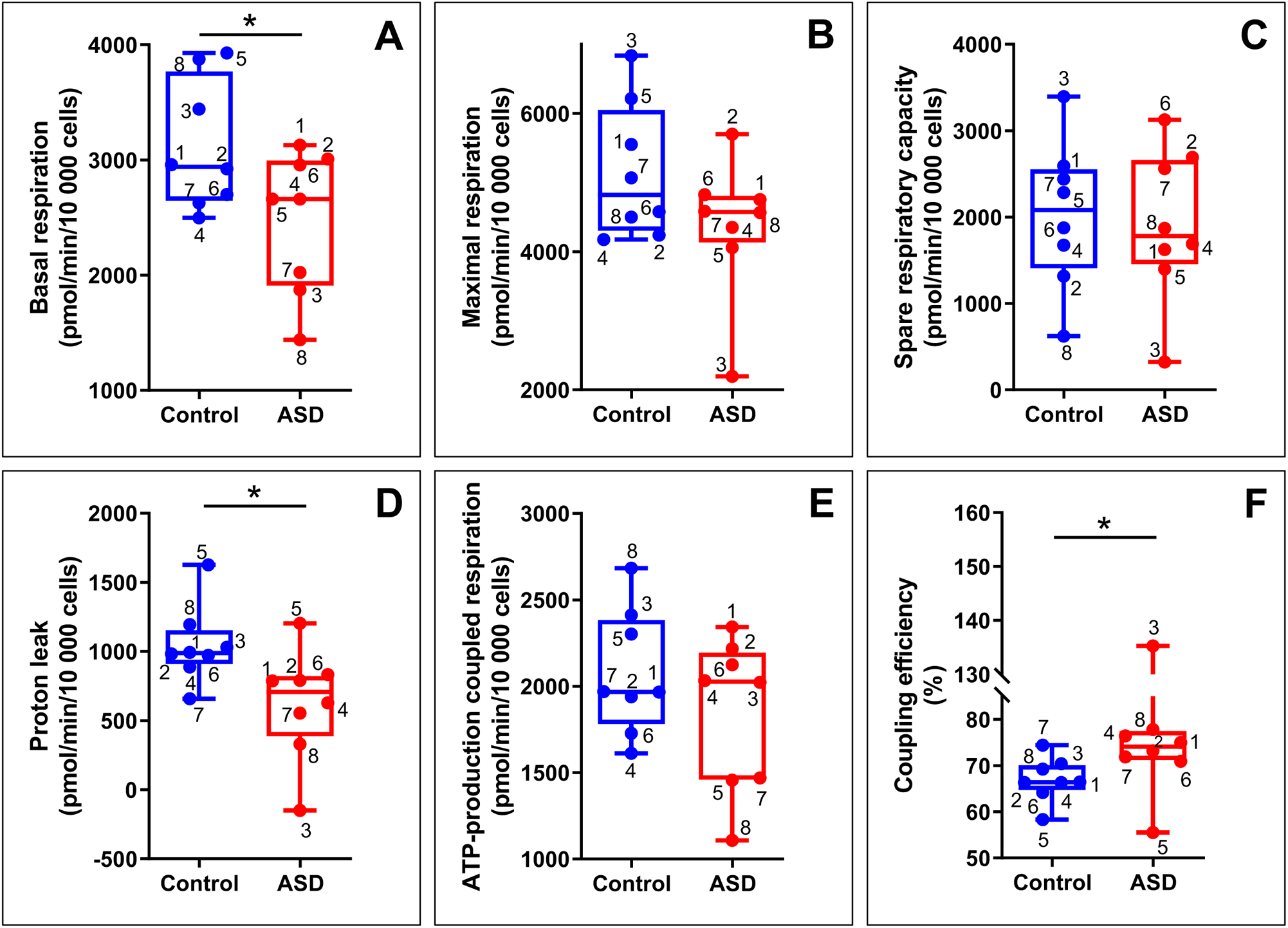
Cell respiration. Stem cells from ASD patients are affected by a decrease in basal respiration (A), proton leak (D) and coupling efficiency (F) but behave like cells from control individuals for maximal respiration (B), spare respiratory capacity (C), ATP-production coupled respiration (E). * = p<0.05

## Discussion

Using stem cells as a mean to assess disturbances arising during the first steps of ontogeny, the current study demonstrates that, when compared with those of control individuals, cells from ASD patients exhibit i) a diminished expression/abundance of glycolysis-associated transcripts and metabolites, ii) an overall reduced ATP production, iii) a lessened basal cell respiration and iv) a modified mitochondrial network. These results confirm and partially invalidate previously published research.

### Transcriptome and metabolome both indicate a reduced glycolysis

Anomalies in the carbohydrate metabolism was described as early as 1985 when a publication described a co-occurrence of lactic acidosis and autism spectrum disorders in 4 young patients (Coleman and Blass 1985). Interestingly, the excess of lactic acid was also associated in one out of four individuals with hyperuricosuria and hyperuricemia. Subsequently, numerous studies reported elevated levels of lactate and pyruvate and a disturbed production of enzymes associated with these metabolites in serum, plasma, urine, fibroblasts and brain (Ramirez-Barbieri et al. 2019; Picard, Wallace, and Burelle 2016; László et al. 1994; Arnold et al. 2003; Goh et al. 2014; Haas 2010; Weissman et al. 2008; Filipek et al. 2003; El-Ansary, Ben Bacha, and Al-Ayadhi 2011). However, the rate of variation of each of these potential biomarkers varies greatly from one study to another, ranging from 17% to 77% (László et al. 1994; Arnold et al. 2003). Nevertheless, we still need to understand the discrepancy between the results of all these studies, which indicate an increase in lactate and pyruvate, and those of our study, which report a decrease in pyruvate in all ASD patients and a mixed variation (overabundance in 3 patients and deficiency in 3 others).

It can be conjectured that an excess of pyruvate in biological fluids reflects either excessive secretion or low input of this metabolite by the cells, leading to an intracellular deficit, as suggested in the current study by the misexpression of transcripts coding for SLC16A9 and SLC16A14, two pyruvate transporters. Only one previous study assessed the quantity of intracellular lactate and pyruvate in fibroblasts and found an increase in the lactate/pyruvate ratio (Weissman et al. 2008). The authors do not give the values for each metabolite, but it is conceivable that the increase in this ratio is partly associated with a decrease in pyruvate production. Furthermore, a decrease in pyruvate dehydrogenase activity has been observed in t he prefrontal cortex of ASD patients (Gu et al. 2013), leading us to imagine either a reduced abundance of upstream pyruvate or a mutation in the gene that would affect the protein’s functionality.

At the transcriptome level, we observed an under-expression of many glycolysis-related genes all along the pathway. At one end, we noticed a misexpression, usually an under-expression, of at least 2 of 4 glucose transporters in every patient. This finding is consistent with studies describing a link between neurodevelopmental disorders, including ASD, and gene anomalies, namely variants or deletions (Mellone et al. 2022; Mir et al. 2022; López-Rivera et al. 2020; Lee, Smith, and Paciorkowski 2015; Redin et al. 2014; Srour et al. 2017). However, at the other end, no previous study reported an association between ASD and molecular anomalies for SLC16A9 and SLC16A14, two pyruvate transporters.

Between these two ends, a misexpression of 6 of the 9 transcripts encoding glycolysis-associated proteins was detected in our ASD cohort. Confirmation can be found in previous studies describing a reduced production of the protein hexokinase 2 (Meyyazhagan et al. 2020) but, more importantly, enolase 2, whose expression is altered in all patients in our cohort, and which is considered by some researchers to be a potential biomarker of ASD (Stancioiu, Bogdan, and Dumitrescu 2023; Ramirez-Celis et al. 2020). *ENO2* hypermethylation was first found in peripheral blood cells from a cohort of 131 age- and sex-matched pairs of ASD subjects and controls in 15% of ASD patients. This gene alteration led to a reduced abundance (70% and 50%, respectively) of the transcript and protein (Wang et al. 2014). Interestingly, this lower intracellular production of enolase is mirrored by a higher abundance in the serum of ASD neonates (Lv et al. 2016) and ASD children (Ayaydın et al. 2020; Stancioiu, Bogdan, and Dumitrescu 2023; Ramirez-Celis et al. 2020). However, it must be noted that one study failed to find a significant difference in serum ENO2 between ASD patients and healthy individuals (Esnafoglu et al. 2017).

### A disrupted biosynthesis of carnitine

As described above, we observed a discrepancy between the lesser abundance of α-ketoglutarate within ASD olfactory stem cells and the upper amount of the same metabolite in a body fluid, namely urine (Harutyunyan, Harutyunyan, and Yenkoyan 2021) of ASD patients. Once again, it may be surmised that these divergent results are the consequence of uncontrolled flux of the molecule out of the cell. In any case, according to our metabolomic data, it seems that the abnormal production of this acid is predominantly associated with the carnitine biosynthesis pathway and not with the citric acid cycle (Krebs cycle). In fact, stem cells of all ASD patients show a disturbed abundance of at least 4 of the 7 measured carnitine-related molecules. Apart from ketoglutarate and ascorbic acid, which are always less abundant, the other 5 metabolites display a variable balance between higher and lower quantities.

As reported in a review (Demarquoy and Demarquoy 2019), reduced serum total and free carnitine has been observed in ASD patients (Filipek et al. 2004), an important finding since this molecule participates to the transport of long-chain fatty acids from the cytosol into the mitochondria. In a large cohort of 213 individuals, 17% of ASD children were found to have elevated in short-chain and long-chain acyl-carnitines (Frye et al. 2013). In addition, the imbalance in carnitine biosynthesis is supported by several genetic anomalies. One team, using two different technical approaches, unveiled exonic copy number variants in the genome of ASD patients and identified in a male proband a deletion of TMLHE (trimethyllysine hydroxylase epsilon), the gene that codes for the first enzyme in the biosynthesis of carnitine, located in mitochondria (Patricia B.S. Celestino-Soper et al. 2011; Patrícia B. S. Celestino-Soper et al. 2012). Homozygous mutations in the gene encoding SLC7A5, an amino acid transporter capable of transporting carnitine, have also been observed in several ASD patients (Tarlungeanu et al. 2016).

A proven carnitine deficiency in some patients suggests that certain symptoms could be improved by supplementation. This was tested in three clinical trials in ASD patients with unknown genetic causes and non-syndromic forms of the disease (for a review, (Malaguarnera and Cauli 2019)). Two assessment scales - CARS (childhood autism rating scale) and CGIS (clinical global impression scale) - were mainly used to evaluate the course of the disorders. The first trial, randomized placebo-controlled with a dose of 50 mg/kg/day, lasted 3 months and demonstrated an improvement in symptoms, according to the CARS and CGIS, correlated with the level of carnitine in the serum (Geier et al. 2011). The second trial, also randomized placebo-controlled but with a dose of 100 mg/kg/day, lasted 6 months and reported an improvement in symptoms according to the CARS criteria but did not observe a link with circulating carnitine levels (Fahmy et al, 2013; Fahmy et al, 2013). The last trial, an open label trial with a dose of 200 mg/kg/day, lasted 8 weeks and observed an improvement according to the CGIS scale (Goin-Kochel et al. 2019).

### A lower abundance of ATP

Our metabolomic data indicate a deficiency of adenine and adenosine diphosphate (ADP) in 8 and 6 ASD patients, respectively. No result being available for adenosine triphosphate (ATP), we measured its production with the Seahorse equipment and protocols. As highly proliferating cells and similarly to cancer cells, stem cells rely primarily on glycolysis, instead of oxidative phosphorylation (OXPHOS), to produce energy from glucose. This metabolic shift is known as the “Warburg effect” (Warburg 1956) and can be visualized on figure 2C. As posited by the altered glycolysis in ASD stem cells described above, a reduced amount of ATP produced by this pathway was observed, a finding consistent with a study showing that cerebral organoids derived for ASD patients exhibit a predominance of glycolysis over OXPHOS and reduced ATP production (Ilieva and Uchida 2022). Nonetheless, it should be noted that such findings depend on the context and, possibly, the cell type. For example, when faced with acute increases in reactive oxygen species, lymphoblastoid cell lines augment glycolysis and glycolytic reserve (Rose et al. 2017).

To compensate for this lack of energy supplied by glucose metabolism, olfactory stem cells produce more ATP through oxidative phosphorylation. However, as this cell type relies heavily on glycolysis for energy, overall ATP production remains deficient. In other cell types, the picture is different. Reduced activity of at least 3 of the 5 respiratory complexes has been observed in muscle cells (Filiano et al. 2002; Weissman et al. 2008; Shoffner et al. 2010) and in lymphoblasts (Giulivi et al. 2010) whereas only abnormalities of complex I have been reported in plasma (Khemakhem et al. 2017). To explain these discrepancies, the time factor must also be considered. For example, diminished activities of various respiratory complexes were found in different brain areas – cerebellum, frontal, parietal, occipital and temporal cortexes – of young children (4-10 years of age) but not in adults (14-39) (Chauhan et al. 2011). The ability of olfactory stem cells to partially remedy their ATP deficit associated with defective glycolysis results from their remarkable mitoplasticity. Thanks to special mitochondrial dynamics, favourable to fission and unfavourable to fusion, cells from ASD patients increased their mitochondrial area and enhanced their ability to produce energy *via* the electron transport chain. Unsurprisingly, our transcriptomic data indicate an exacerbated expression of genes encoded by mitochondria in ASD cells, with a special mention for NT-ND6 whose expression is on average doubled when compared with control cells (data not shown). Similarly, smaller more fragmented mitochondria were observed in the black and tan, brachyury (BTBR) mouse model of ASD (Ahn et al. 2020). Atypical mitochondria morphology was also observed in ASD fibroblasts (Pecorelli et al. 2020). In other words, mitoplasticity may represent one of the compensatory mechanisms by which ASD cells attempt to adapt to altered energy supply and need.

### A diminished basal respiration

A relatively small number of studies investigated cellular respiration in ASD patients. It was first demonstrated that, at baseline and after exposure to reactive oxygen species, lymphoblastoid cell lines from some children with ASD increase ATP-linked respiration as well as maximal and reserve respiratory capacity (Rose et al. 2014; 2015; 2017). The same research group, comparing serum microRNAs between healthy individuals and ASD patients, observed negative correlations between miRNA levels and mitochondrial respiration in a subgroup of patients (Jyonouchi et al. 2019). In agreement with the former findings, higher respiratory capacities and complex IV activity were reported in a lymphoblastoid cell line derived from an autistic child (Hassan et al. 2021). Finally, peripheral blood mononuclear cells indicate that patients with ASD have increased respiratory reserve capacity and maximal respiration (Gevezova et al. 2021).

None of these results are consistent with the conclusions of this study. Maximal respiration and spare respiratory capacity are not exacerbated while basal respiration and proton leak are reduced. The cause of these anomalies is not the consequence of a defective expression of gene families – NDUFs (complex I), SDHs (complex II), UQCRs (complex III), COXs (complex IV), ATP5s (complex V) – associated to oxidative phosphorylation since, according to our transcriptomic data, no difference was observed between ASD patients and control individuals. However, more than the quantity of proteins, it is perhaps their quality that is at stake in this metabolic pathway. Sequencing of the genomes of the 8 autistic patients (data not shown) indicates that they suffer from exonic frameshift deletions in several genes coding for proteins involved in oxidative phosphorylation. Of note, the cells of patients 3, 7 and 8 that display the most altered basal respiration are also those that are affected by the most numerous exonic frameshift deletions – 9, 10 and 10 respectively - in complexes IV and V. However, in the absence of a precise assessment of the functional consequences of these deletions, all this remains highly speculative.

### Limits of the study

For the current study and two previous ones, we analyzed stem cells in order to be as close as possible to the first stages of development. However, these cells are highly proliferative and display an unalike metabolism to differentiated cells. This has positive aspects. For example, due to a particularly high level of glycolysis, we were able to demonstrate a defect in ATP production associated with this metabolic pathway. However, this phenomenon should be marginal in cells in which glycolytic ATP represents 7% rather than 70% of the total production of energy molecules.

The reasons for this disturbed glycolysis probably need to be sought at the genomic level. Following sequencing of the genome of stem cells from the cohort of autistic patients, we looked for a possible mutation in the exons of the gene coding for enolase 2, a gene whose transcription is strongly disrupted in all ASD patients. However, we did not observe any such mutation. We therefore need to look for a possible methylation defect in the promoter region of this gene. This experiment is all the more necessary as a study has shown such a phenomenon in peripheral blood cells (Wang et al. 2014).

Although easily accessible in a healthy, compliant subject, nasal olfactory stem cells are not in ASD patients. To confirm our results and possibly use them as biomarkers, it is important to conduct a similar study on peripheral blood cells with a larger cohort of individuals. It would also be interesting to compare the two sexes in order to study any discrepancy.

## Acknowledgements

High throughput sequencing was performed at the TGML Platform, supported by grants from Inserm, GIS IBiSA, Aix-Marseille Université, and ANR-10-INBS-0009-10. We are grateful to Thi Tien Nguyen and Alice Carrier for their help and insight on Seahorse data. The work was funded by CNRS (ref MOTISM), Aix-Marseille Université and Fondation de l’Avenir.

